# Dasatinib Resensitizes MAPK Inhibitor Efficacy in Standard-of-Care Relapsed Melanomas

**DOI:** 10.1101/2023.01.20.524923

**Authors:** Vito W. Rebecca, Min Xiao, Andrew Kossenkov, Tetiana Godok, Gregory Schuyler Brown, Dylan Fingerman, Gretchen M. Alicea, Meihan Wei, Hongkai Ji, Jeremy Bravo, Yeqing Chen, Mitchell E. Fane, Jessie Villanueva, Katherine Nathanson, Qin Liu, Y. N. Vashisht Gopal, Michael A. Davies, Meenhard Herlyn

## Abstract

Resistance to combination BRAF/MEK inhibitor (BRAFi/MEKi) therapy arises in nearly every patient with *BRAF*^V600E/K^ melanoma, despite promising initial responses. Achieving cures in this expanding BRAFi/MEKi-resistant cohort represents one of the greatest challenges to the field; few experience additional durable benefit from immunotherapy and no alternative therapies exist. To better personalize therapy in cancer patients to address therapy relapse, umbrella trials have been initiated whereby genomic sequencing of a panel of potentially actionable targets guide therapy selection for patients; however, the superior efficacy of such approaches remains to be seen. We here test the robustness of the umbrella trial rationale by analyzing relationships between genomic status of a gene and the downstream consequences at the protein level of related pathway, which find poor relationships between mutations, copy number amplification, and protein level. To profile candidate therapeutic strategies that may offer clinical benefit in the context of acquired BRAFi/MEKi resistance, we established a repository of <u>p</u>atient-<u>d</u>erived <u>x</u>enograft models from heavily pretreated patients with resistance to BRAFi/MEKi and/or immunotherapy (R-PDX). With these R-PDXs, we executed *in vivo* compound repurposing screens using 11 FDA-approved agents from an NCI-portfolio with pan-RTK, non-RTK and/or PI3K-mTOR specificity. We identify dasatinib as capable of restoring BRAFi/MEKi antitumor efficacy in ∼70% of R-PDX tested. A systems-biology analysis indicates elevated baseline protein expression of canonical drivers of therapy resistance (e.g., AXL, YAP, HSP70, phospho-AKT) as predictive of MAPKi/dasatinib sensitivity. We therefore propose that dasatinib-based MAPKi therapy may restore antitumor efficacy in patients that have relapsed to standard-of-care therapy by broadly targeting proteins critical in melanoma therapy escape. Further, we submit that this experimental PDX paradigm could potentially improve preclinical evaluation of therapeutic modalities and augment our ability to identify biomarker-defined patient subsets that may respond to a given clinical trial.

**SINGLE SENTENCE SUMMARY:** Broad target inhibition effective as a salvage strategy in BRAF/MEK inhibitor-acquired resistance PDX

## INTRODUCTION

There has been immense progress in the available treatment for patients with metastatic melanoma, with 14 therapies FDA-approved since 2011. Vertical targeting of the mitogen-activated protein kinase (MAPK) pathway with the use of dual BRAF/MEK inhibitor (BRAFi/MEKi) therapy for patients with *BRAF*^V600E/K^ mutant melanoma, as well as immune checkpoint inhibitor-based strategies available for all genotypes of melanoma have each improved the overall survival of patients with advanced disease. However, long-term responses beyond 5 years are only seen in 25-35% of patients, due to the persistence of therapy-resistant subpopulations of melanoma cells that employ diverse mechanisms of escape to drive intrinsic and acquired resistance, including cell state- and signaling pathway-plasticity. In the way of cell state plasticity that enable therapy escape, melanoma cells can dedifferentiate to occupy a variety of alternative identities including a neural crest stem cell-like identity (1), an invasive identity, and a stress cell identity (2). Regarding signaling pathway plasticity, a growing body of evidence identifies hyperactivation of compensatory signaling through the PI3K/AKT/mTORC1 (3), SRC (4), and STAT3 (5) pathways via upregulation of receptor tyrosine kinases (RTKs) including insulin growth factor receptor 1 (IGFR-1) (6), AXL (7), PDGFRβ (8), EGFR (9), c-MET (10), and HER3 (ERBB3) (11). However, efforts to target these diverse resistance mechanisms have not resulted in a novel approved-therapy regimen to address patients who have already relapsed to standard-of-care (SOC) therapies.

Due to the immense tumoral heterogeneity across melanoma patients and the need for more personalized therapy strategies, “master” protocols have been created to provide medications better tailored to the unique genetic makeup of a given patient’s tumor (12). The national cancer institute (NCI) is currently testing a master protocol in the NCI molecular analysis for therapy choice (MATCH) trial, whereby cancer patients are provided therapy regimens based on the genetic changes found in their tumors (13, 14). Although highly innovative and geared to better address tumor heterogeneity, it remains to be seen how effective this strategy will be. Trials such as the NCI-MATCH are built upon a set of scientific assumptions including a) a correlation between an activating mutation in a pathway node and elevated activation of that respective pathway (e.g., *PI3K* mutations correlate with activation of PI3K/AKT/mTOR at the protein level), b) copy number increases of a gene correlate with the protein expression of that gene (e.g., tumors with *EGFR* amplification would exhibit increased EGFR protein expression), and c) tumors that harbor a given mutation are more sensitive to inhibitors that target the related pathway (e.g., tumors that harbor *AKT* mutations should be more sensitive to an mTOR inhibitor). There are genomic alterations that predict sensitivity to certain compounds, including *BRAF*^V600E/K^ mutations for V600E/K selective BRAF inhibitors (e.g., vemurafenib) in melanoma patients and *EGFR* mutations for EGFR inhibitors (e.g., gefitinib) in patients with advanced lung adenocarcinoma, however how robust the relationship between genomic status and antitumor efficacy of a cognate inhibitor is remains to be fully understood.

Here, we aim to develop a therapy regimen for melanoma patients that have already relapsed to SOC therapies and for which no effective therapy regimens remain. Using a genomically, transcriptionally, and proteomically characterized set of patient-derived xenograft (PDX) models of metastatic melanoma, the relationship between mutation, copy number status, and total-/phospho-protein expression was analyzed, revealing a poor correlation. These analyses were expanded to cutaneous melanoma patient samples in the TCGA where reverse-phase protein array, copy number variation, and mutational information is available, which also corroborated the poor relationship between genomic and protein status. To develop an effective salvage therapy strategy, we developed a PDX therapy repurposing screen to investigate the potential for a triple combination including BRAFi/MEKi and a third inhibitor against the PI3K/AKT/mTOR pathway or a pan-RTK inhibitors to increase the overall survival relative to BRAFi/MEKi treatment alone. Relationships between gene and protein status with therapy efficacy reveal benefit in assessing total- and phospho-protein expression when determining tailored therapy strategies for patients. Of the 11 triple combinations screened, the most effective consisted of BRAFi/MEKi plus dasatinib. Sensitivity analysis and scRNAseq suggest the robust efficacy of the dasatinib combination in 14 models of BRAFi/MEKi-resistant PDX models relies on the broad target profile of dasatinib capable of suppressing diverse melanoma subpopulations critical in minimal residual disease and therapy escape.

## RESULTS

### BRAFi/MEKi resistant tumors display elevated RTK/MAPK/PI3K/mTOR activity

The majority of patients with advanced *BRAF*^V600E/K^ mutant melanoma will progress while on BRAFi/MEKi therapy. Further, four out of five of these patients show no long-term benefit from immunotherapy (i.e., anti-PD-1, anti-CTLA-4) due to diverse mechanisms that confer cross-resistance across different therapeutic modalities (15). Therefore, there is an unmet need for durable second- and/or third-line (salvage) therapy strategies for patients that have already relapsed on standard-of-care (SOC) strategies. Although the list of reported resistance mechanisms to BRAFi/MEKi continues to grow (16-23), they predominately center around reactivation of the MAPK pathway and the hyperactivation of parallel survival signaling through the PI3K/AKT/mTOR pathway via elevated receptor tyrosine kinase (RTKs) and non-receptor kinase activity.

In agreement, reverse-phase protein array (RPPA) analyses of 94 treatment-naïve, 22 BRAFi-resistant, and 16 BRAFi/MEKi-resistant patient-derived xenograft (PDX) models reveal a heterogeneous enrichment in various total and phospho-proteins involved in RTK, PI3K/AKT/mTOR, and MAPK signaling (**Figure 1A**). IGFBP2, phospho-HER3 (Y1289), phospho-cMET (Y1235), phospho-MEK (S217/S221), and phospho- ERK1/2 (T202/Y204) levels were significantly elevated amongst BRAFi- and BRAFi/MEKi-resistant PDX models relative to therapy naïve models. AXL, a dogmatic marker of a therapy resistance melanoma cell state was not significantly increased in BRAFi- and BRAFi/MEKi-resistant models. However, reduced MITF expression was observed, suggesting melanoma models with acquired resistance occupy a de-differentiated cell state, as been previously suggested. Of note, multiple nodes of the PI3K/AKT/mTOR pathway (e.g., phospho-AKT T308/S473, phospho-mTOR S2448, phospho-4E-BP1 T37/T46/S65, phospho-S6 S235/S236/S240/S244) were not significantly different between therapy resistant PDX models and therapy-naïve models (**Figure 1A**).

**FIGURE 1:**
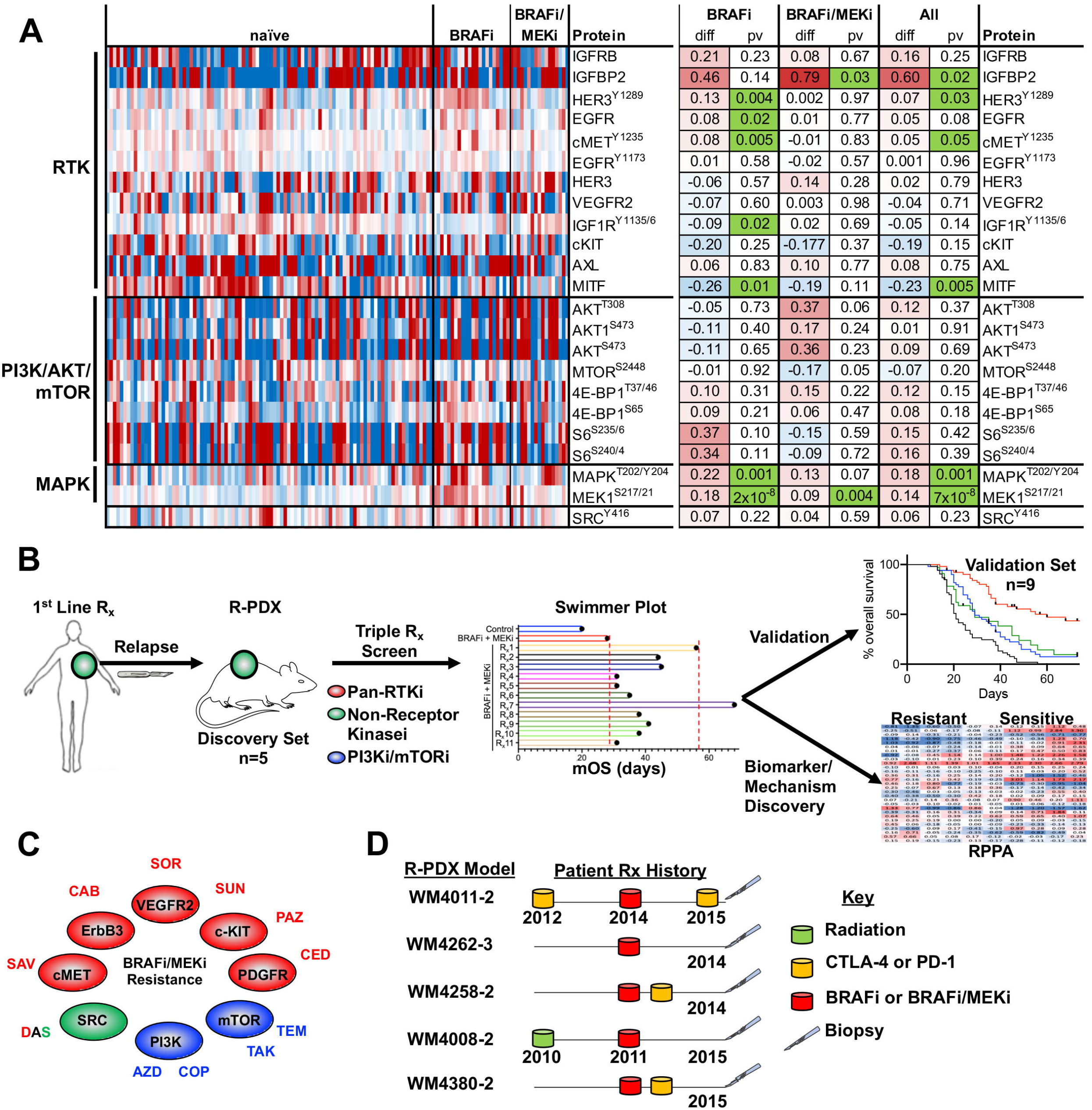
BRAFi/MEKi resistant tumors display elevated RTK/MAPK/PI3K/mTOR activity. (**A**) Differential total- and phospho-protein expression analysis between therapy naïve PDX, BRAFi-R PDX and BRAFi/MEKi-R PDX models. (**B**) Graphical summary of the experimental strategy of this manuscript. (**C**) Targets of the 11 FDA-approved compounds used in our repurposing screen. (**D**) Therapy history of the 5 patients whose tumors were used to establish R-PDX that were subsequently screened against 11 BRAFi/MEKi-based triple combinations.

To begin our efforts to develop an effective salvage strategy for patients that have relapsed SOC therapy, we designed a custom drug screen using 11 Food and Drug Administration (FDA)-approved compounds from a National Cancer Institute (NCI) drug repository that possess pan-RTK, non-RTK and/or PI3K-mTOR specificity (**Figure 1B, 1C, Table 1**). Each of the compounds used in this study have been previously tested in advanced melanoma patients, however not in combination with BRAFi/MEKi and not in the context of patients who have previously relapsed to BRAFi or BRAFi/MEKi in a stage II or greater trial. With the goal to identify a three-drug cocktail consisting of BRAFi/MEKi in combination with a third compound that possesses robust second- and/or third-line efficacy, we initially selected a discovery set of 5 R-PDX models derived from patients that have previously progressed while on treatment with BRAFi in the presence or absence of MEKi, 3 of these patients also progressed while receiving immune checkpoint blockade (anti-CTLA-4 or anti-PD-1) and 1 patient also progressed after receiving radiation therapy to identify an effective triple combination we would then validate in an expanded cohort of R-PDX (**Figure 1D**).

### Protein expression nor genetic status of the PI3K/-mTOR pathway robustly predicts efficacy of PI3K/mTOR pathway inhibitors

The clinical community has begun using *umbrella trial* approaches to better personalize therapy, whereby a “master protocol” is designed to evaluate multiple investigational drugs based off of characterization of tumor DNA from a given patient via targeted sequencing to determine potential therapy combinations that incorporate an inhibitor paired against a related mutation unique to that patient’s tumor (12, 24). The assumptions built into the umbrella trial strategy include the 1) gene mutational status correlates with activation at the protein level of the associated pathway, and 2) gene mutational status correlates with tumor sensitivity towards a cognate inhibitor (**Figure 2A**). For example, a patient whose tumor harbors an *AKT* mutation is expected to a) possess elevated AKT pathway activity at the protein level (e.g., elevated phospho-AKT S473) and b) be relatively more sensitive to an AKT or downstream mTOR inhibitor relative to a patient with wild type *AKT*. We tested the robustness of the umbrella trial strategy in predicting sensitivity to 4 PI3K-mTOR pathway inhibitors included within our repurposing screen amongst the 5 discovery R-PDX models. We performed targeted sequencing and copy number variation (CNV) analyses of genes in the PI3K/AKT axis (**Figure 2B**), as well as reverse phase protein arrays to determine the baseline activation status of the PI3K/mTOR pathway in our discovery set of R-PDX models (**Figure 2C**). We observed a) no correlation between AKT3 copy number amplification and AKT activity at the protein level, and b) no correlation between *NRAS* mutation and hyperactivation of PI3K/AKT and MAPK signaling at the protein level. In agreement with previous findings, *PTEN* copy number loss was associated with the greatest hyperactivation of PI3K/AKT signaling, as seen in the WM4008 model.

**FIGURE 2:**
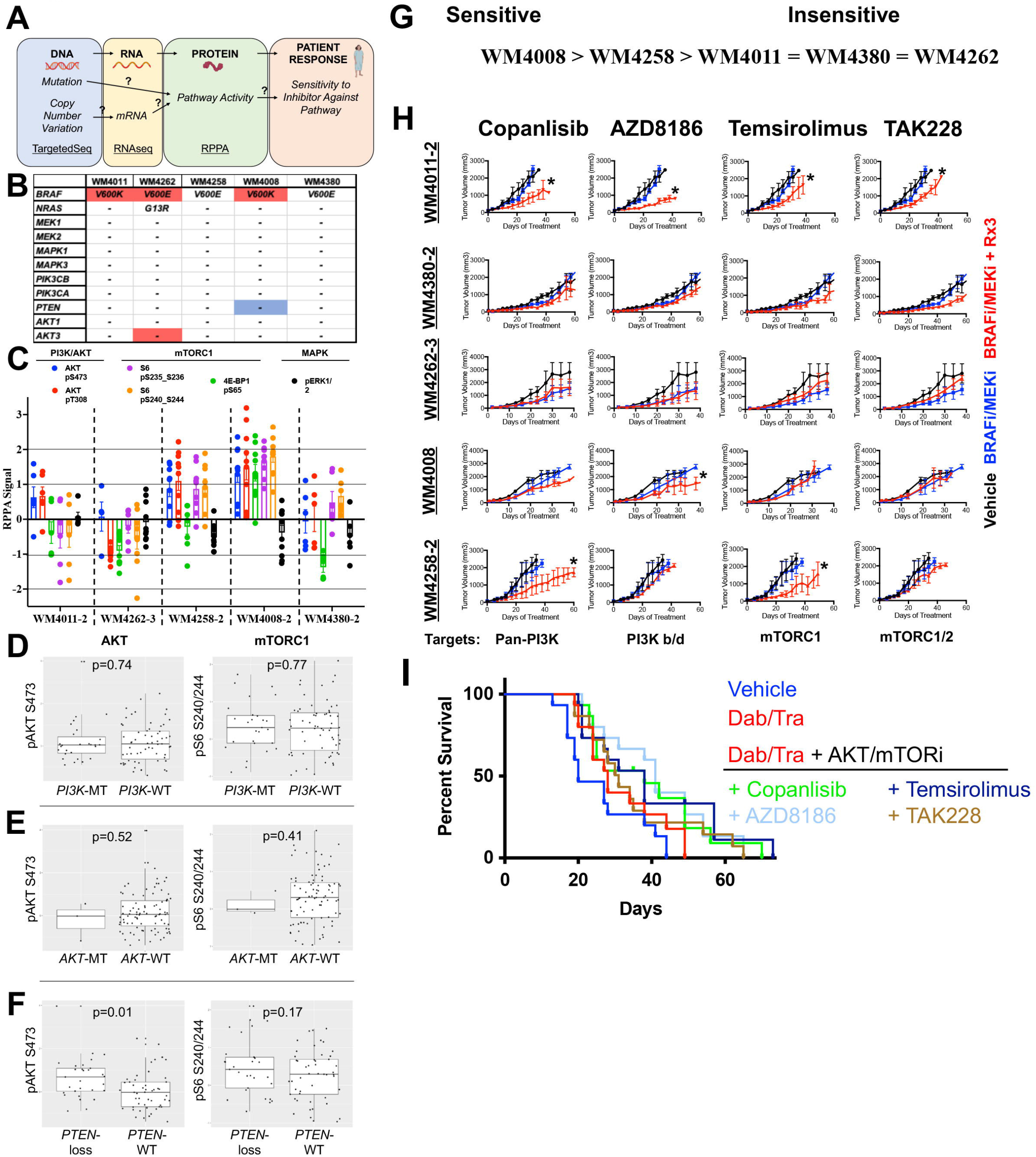
Protein expression nor genetic status of the PI3K/-mTOR pathway robustly predict efficacy of PI3K/mTOR pathway inhibitors. (**A**) Schematic for the role of the central dogma theory in clinical practice. “?”s denote assumptions with poor validation in the melanoma literature. (**B**) Mutational and CNV analysis of PI3K/AKT pathway nodes across the R-PDX models in the discovery screen. (**C**) RPPA analysis of PI3K/AKT pathway nodes across the R-PDX models in the discovery screen. (**D**) Analysis of the relationship between AKT/mTOR protein activation status and *PI3K* mutations (**E**) *AKT* mutations, and (**F**) *PTEN* copy number loss in cutaneous melanoma patients in the TCGA. (**G**) Therapy sensitivity prediction analysis based on the unique genomic and/or proteomic signature of each R-PDX and the protein targets of a given agent. (**H**) Tumor volumes for NSG mice implanted with 1 of the 5 R-PDX organized by row and treated with vehicle (black line), BRAFi/MEKi (blue line), or BRAFi/MEKi plus a third agent (red line) as labeled. (**I**) Overall survival plots of R-PDX bearing mice treated with the therapies as shown. * signifies p<0.05.

Given the small sample size amongst our R-PDX, we analyzed the relationship between genomic status and pathway activation in 90 melanoma patients in the TCGA where RPPA and genomic information is available. Notably, patients whose tumors harbor *PIK3C* mutations did not display elevated AKT (pAKT S473) or mTORC1 (pS6 S240/244) activity at the protein level relative to patients whose tumors are *PIK3C* wild type (**Figure 2D**). Further, patients whose tumor harbor *AKT* mutations also did not display elevated AKT or mTORC1 activity at the protein level relative to patients whose tumors are *AKT* wild type (**Figure 2E**). Patient *PTEN* status again inversely correlated with AKT activity, however downstream mTORC1 activity did not correlate with *PTEN* status (**Figure 2F**).

Based off our multi-omic characterization, we classified R-PDX in our discovery set as potentially sensitive or insensitive to PI3K or mTOR inhibition, whereby WM4008 was predicted to be the most sensitive (based on *PTEN* copy number loss and elevated phospho-AKT and phosho-S6 levels) and WM4262 was predicted to be the most insensitive (based on low phospho-AKT and phospho-S6 levels) (**Figure 2G**). To begin our therapy trials, we again confirmed that all 5 of the discovery set R-PDX models were resistant to BRAFi/MEKi, as seen in the near identical tumor growth patterns in tumor bearing NSG mice treated with vehicle control versus a combination of BRAFi (dabrafenib) and MEKi (trametinib) (**Figure 2H, 2I**). Treatment with the pan-PI3K inhibitor copanlisib in combination with BRAFi/MEKi conferred a significant antitumor response in 2 of the 5 R-PDX relative to BRAFi/MEKi treatment. A poor relationship between PI3K/AKT pathway status and copanlisib + BRAFi/MEKi efficacy was observed, whereby the R-PDX (WM4008) predicted to be most sensitive did not experience a response. Treatment with the PI3K β/δ isoform specific inhibitor AZD8186 also elicited a significant antitumor response in 2 of the 5 R-PDX relative to BRAFi/MEKi treatment. In this case, AZD8186 modestly increases the antitumor activity of BRAFi/MEKi in the R-PDX predicted to be sensitive to a PI3K/AKT pathway inhibitor (**Figure 2G, 2H**). We next tested the efficacy of the allosteric and catalytic mTOR inhibitors Temsirolimus and TAK228, respectively. Temsirolimus + BRAFi/MEKi modestly increased the antitumor activity of BRAFi/MEKi in 2 of the 5 R-PDX relative to BRAFi/MEKi treatment, however the R-PDX predicted to be sensitive (highest mTOR activity as seen by high phospho-S6 and phospho-4E-BP1 levels) did not respond. Treatment with TAK228 + BRAFi/MEKi demonstrated mild antitumor activity in 1 of the 5 R-PDX, and again did not have efficacy in the R-PDX predicted to be most sensitive. Altogether, these data from our small discovery panel of R-PDX models suggest that treatment decisions for patients enrolled in basket trials with PI3K/mTOR inhibitors may not be optimal if solely based on genetic, genomic, or proteomic information of the PI3K/mTOR pathway. Further, the genetic status of the PI3K/AKT pathway is not predictive of AKT and mTORC1 activity at the protein level, suggesting better genetic biomarkers of PI3K/AKT pathway activity are needed.

### Protein expression, not genetic status, of RTKs partially predicts efficacy of pan-RTK inhibitors

We next tested the robustness of the umbrella trial approach in predicting R-PDX sensitivity to the 6 RTK inhibitors in our repurposing screen (**Figure 1C**). Mutational and CNV analysis of a panel of RTK genes in our targeted sequencing panel revealed copy number amplifications in *EGFR* and *MET* in 1 and 2 R-PDX models, respectively (**Figure 3A**). In addition, at least 1 mutation in *EGFR, ERBB2, ERBB4, FGF4*, and/or *FGFR3* was detectable in 4 of the 5 R-PDX. As could be expected, RPPA analyses revealed heterogenous RTK protein expression and activation status across the discovery set of R-PDX models (**Figure 3B**). Notably, there was poor relationship between CNV status and protein expression of a given RTK. For example, the R-PDX WM4262 possesses *EGFR* copy number amplification, however, does not display elevated EGFR protein expression relative to R-PDX without *EGFR* copy number gains. Given the small sample size amongst our R-PDX, we again analyzed the relationship between genomic status and protein expression of RTKs in 90 melanoma patients in the TCGA where RPPA and genomic information is available (**Figure 3C, D, R, F, G, H**). Notably, there was no significant correlation between copy number status and protein expression of *EGFR, PDGFR*β, *HER2, VEGFR2, HER3* and *AXL* between patients with gene amplifications and patients with normal copy number.

**FIGURE 3:**
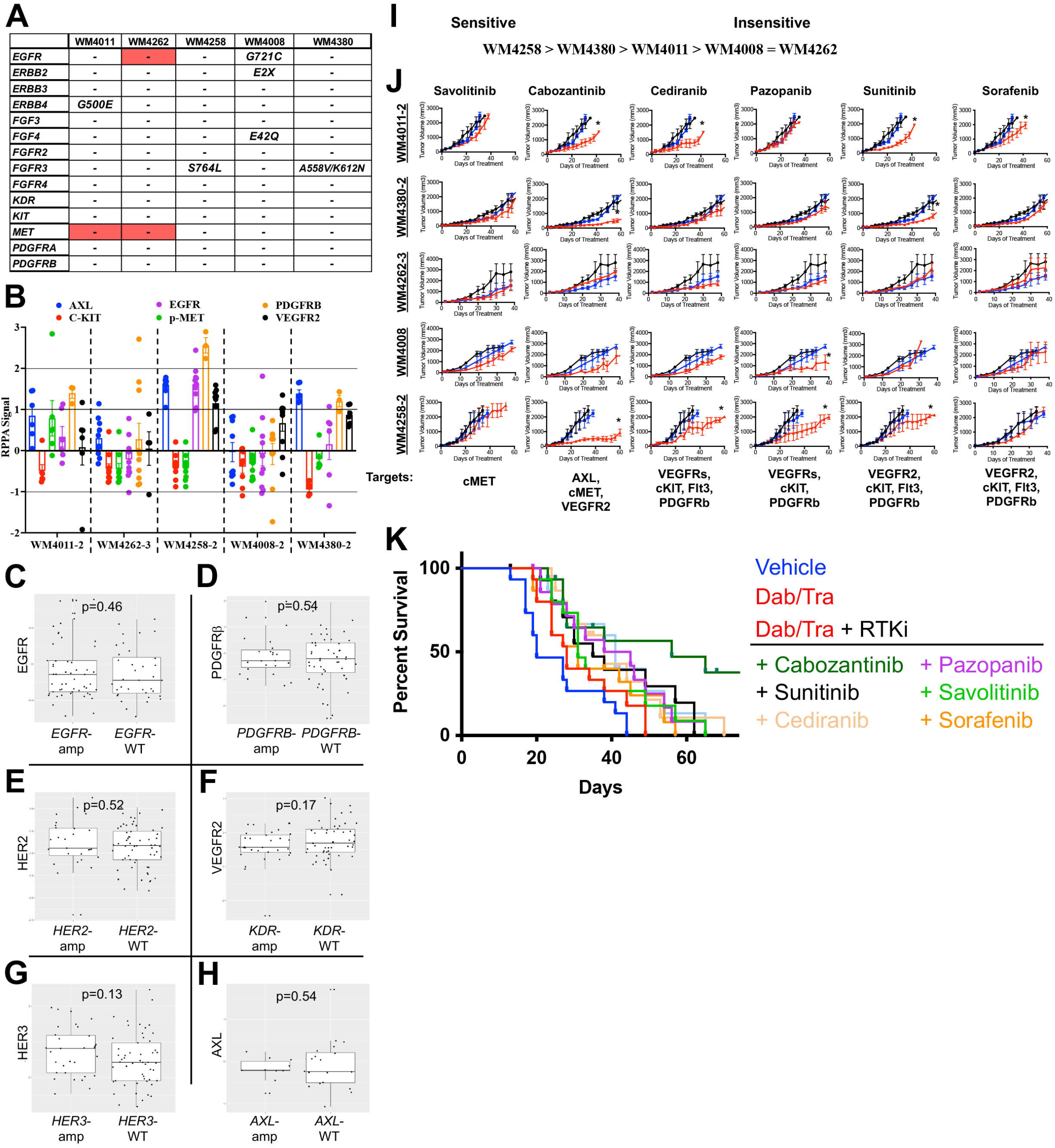
Protein expression, not genetic status, of RTKs partially predicts efficacy of pan-RTK inhibitors. (**A**) Mutational and CNV analysis of RTK nodes across the R-PDX models in the discovery screen. (**B**) RPPA analysis of RTK nodes across the R-PDX models in the discovery screen. (**C**) Analysis of the relationship between RTK protein expression and gene copy number gains in *EGFR*, (**D**) *PDGFRβ*, (**E**) *HER2*, (**F**) *VEGFR2*, (**G**) *HER3*, (**H**) *AXL*. (**I**) Therapy sensitivity prediction analysis based on the unique genomic and/or proteomic signature of each R-PDX and the protein targets of a given agent. (**J**) Tumor volumes for NSG mice implanted with 1 of the 5 R-PDX organized by row and treated with vehicle (black line), BRAFi/MEKi (blue line), or BRAFi/MEKi plus a third agent (red line) as labeled. (**K**) Overall survival plots of R-PDX bearing mice treated with the therapies as shown. * signifies p<0.05.

Based off this genomic and proteomic characterization, we classified R-PDX in our discovery set as potentially sensitive or insensitive to pan-RTK inhibition, WM4258 predicted to be the most sensitive (due to greatest pan-RTK expression) and WM4008 and WM4262 predicted to be least sensitive (**Figure 3I**). Of note, phospho-cMET levels did not vary significantly across the R-PDX models, therefore differences in sensitivity to cMET inhibition were not predicted to occur. Treatment with the selective cMET inhibitor savolitinib + BRAFi/MEKi did not confer any antitumor benefit across the discovery R-PDX panel (**Figure 3J, 3K**). Treatment with the pan-RTK inhibitor cabozantinib (XL-184) + BRAFi/MEKi conferred antitumor activity in 3 of the 5 R-PDX. Notably, the cabozantinib cocktail possessed the antitumor activity in the 3 R-PDX predicted to have sensitivity, based on their RTK protein expression (e.g., PDGFRβ). The pan-RTK inhibitor cediranib revealed a similar trend, conferring antitumor activity in 2 of the 5 R-PDX, both of which were predicted to be sensitive to pan-RTK inhibition. To a lesser extent, pazopanib and sunitinib also demonstrated modest efficacy in R-PDX with elevated RTK protein expression, with the exception of pazopanib + BRAFi/MEKi antitumor activity in 1 R-PDX not predicted to respond due to low RTK expression (WM4008). The exercise of treating our discovery set of R-PDX models with a panel of 6 RTK inhibitors revealed a poor relationship between the mutational or copy number status of a given receptor and the efficacy of its cognate inhibitor. In contrast, an encouraging relationship between RTK protein expression and pan-RTK inhibitor sensitivity emerged. Altogether, these data suggest that treatment decisions for patients enrolled in basket trials would benefit from IHC verification of protein expression of drug targets in addition to genetic/genomic information.

### Dasatinib resensitizes R-PDX models with high expression of canonical melanoma therapy resistance signatures

Of the 11 triple combinations consisting of BRAFi/MEKi plus a third agent, the BRAFi/MEKi plus dasatinib combination emerged as the most effective regimen to increase the mOS of R-PDX bearing mice (**Figure 4A, 4B, 4C**). To test the robustness of the dasatinib + BRAFi/MEKi regimen, we screened an expanded validation set of 9 additional R-PDX models and found the dasatinib + BRAFi/MEKi combination increased mOS of R-PDX bearing mice two-fold relative to mice treated with BRAFi/MEKi or dasatinib alone (**Figure 4D**). Notably, treatment of a therapy naïve PDX model with BRAFi/MEKi + dasatinib did not extent the time to relapse relative to BRAFi/MEKi alone, suggesting this triple combination may better be suited as a second-or third-line therapy to address acquired-rather than intrinsic-resistance mechanisms (**Supplemental Figure 1**). To identify potential biomarkers and understand the underlying mechanism of action, we next interrogated differential total- and phospho-protein expression levels between R-PDX most sensitive and least sensitive to the dasatinib + BRAFi/MEKi cocktail. Notably, R-PDX models most sensitive to the dasatinib + BRAFi/MEKi combination exhibit elevated levels of canonical drivers of therapy resistance in melanoma including HSP70, AXL, phospho-AKT S473, phospho-AKT T308, and YAP. In addition, R-PDX most sensitive to dasatinib _ BRAFi/MEKi displayed elevated protein levels of MRAP, p27, IRS1, UBAC1, LAD1, Notch, XPF, and c-ABL and b) reduced protein levels of PTEN, ATG4B, MYT1, GAB2, and DDB1 (**Figure 4E**). Analysis of markers of sensitivity to the dasatinib + BRAFi/MEKi combination exhibit a negative correlation with the melanocyte lineage factor MITF, suggesting that the dasatinib therapy combination is most effective against de-differentiated melanomas that represent the most difficult to clinically address (**Figure 4F, 4G, 4H, 4I**) (1, 16). Further, analysis of reported dasatinib targets across minimal residual disease (MRD) subpopulations shown to drive therapy resistance in patient scRNAseq data reveal dasatinib targets expressed in distinct subpopulations (**Figure 4J**) (1, 25). For example, the dasatinib target PDGFRβ is highly expressed in neural crest-like stem cell melanoma subpopulations, but lowly expressed in invasive, pigmented and starved-like melanoma cell populations (1). In contrast, AXL is highly expressed in the invasive subpopulation but lowly expressed in other MRD subpopulations. These data suggest dasatinib may have potent activity against R-PDX due to its broad target profile capable of concurrently suppress distinct MRD subpopulations. In a final set of experiments, WM4258 (R-PDX) bearing mice were treated with BRAFi/MEKi plus or minus dasatinib and characterized by scRNAseq, revealing 3 melanoma subpopulations depleted following the addition of dasatinib to BRAFi/MEKi that exhibit a) YAP gene signatures defined by elevated downstream effectors (e.g., CENPF, RRM3), b) dormancy and neural crest stem cell-like markers (e.g., NR2F1, SOX4), RTK nodes (e.g., FGFR1, MET, HGF), and d) and antiapoptotic nodes (e.g., BID; cluster 5) (**Figure 4K, 4L, 4M**). These data suggest the dasatinib + BRAFi/MEKi combination has the potential to address acquired BRAFi/MEKi-resistance due to its broad target profile capable of concurrently eliminating distinct MRD subpopulations.

**FIGURE 4:**
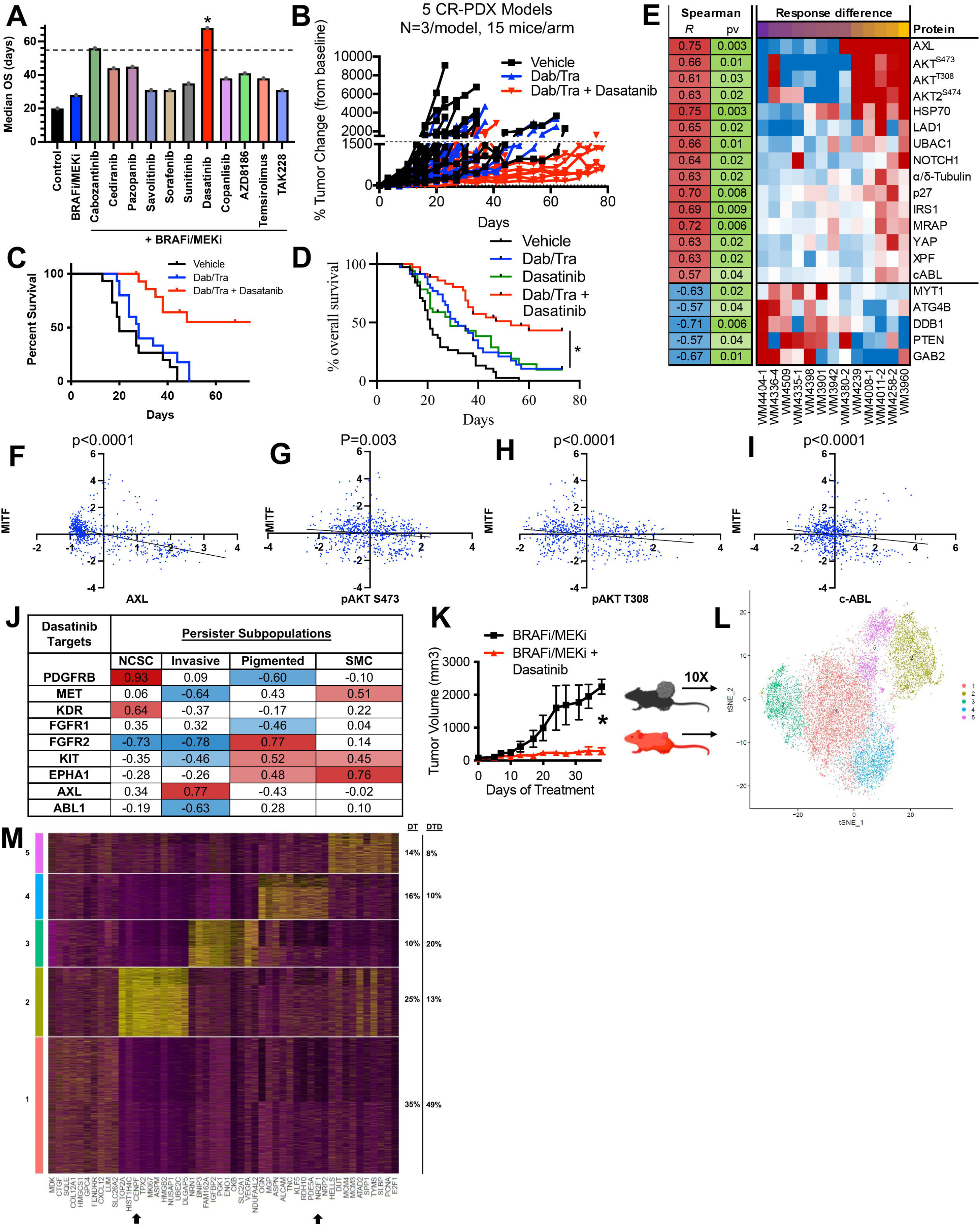
Dasatinib resensitizes R-PDX models with high expression of canonical melanoma therapy resistance mechanisms. (**A**) Waterfall plots depicting median overall survival for the R-PDX treated with the 11 BRAFi/MEKi-based triple combinations. (**B**) Spider plot for discovery set of R-PDX bearing mice treated with vehicle (black line), BRAFi/MEKi (blue line), or BRAFi/MEKi plus dasatinib (red line). (**C**) Overall survival plots of discovery set of R-PDX bearing mice treated with the therapies as shown. (**D**) Overall survival plots of expanded validation set of 9 additional R-PDX bearing mice treated as shown. (**E**) Differential total- and phospho-protein correlation analysis with sensitivity with BRAFi/MEKi plus dasatinib. (**F**) Paired protein expression analysis of MITF with AXL, (**G**) phospho-AKT S473, (**H**) phospho-AKT T308, and (**I**) c-ABL. (**J**) Correlation analysis of dasatinib targets and distinct melanoma populations dominant in minimal residual disease in scRNAseq data from melanoma patients (Tirosh et al, *Science* 2016). (**K**) NSG mice implanted with the R-PDX WM4258 and treated as shown once tumors were palpable. (**L**) t-SNE plot of scRNAseq data from WM4258 tumor cells treated *in vivo* with BRAFi/MEKi plus or minus dasatinib from (K). (**M**) Top gene markers for melanoma subpopulations that are enriched (clusters 1 and 3) or depleted (clusters 2, 4, and 5) following the addition of dasatinib to BRAFi/MEKi. * signifies p<0.05.

## DISCUSSION

We here have demonstrated the limitation of the central dogma theory in the context of metastatic melanoma, with a lack of robust correlations between copy number amplifications and the protein expression of RTKs. Further, mutations in *PI3K/AKT* do not robustly correlate with downstream activation of AKT and mTORC1 at the protein level, which suggests clinical decisions regarding therapy may not be completely founded based off of genomic characterization alone. Although *PTEN* copy number losses correlate with reduced phospho-AKT levels as could be expected, downstream mTORC1 activity did not correlate with *PTEN* loss, which may have implications on whether mTOR inhibitors should be used in this context. Overall, our correlation studies between mutations CNVs, and protein activity suggests IHC analyses of tumor tissue accompany genetic characterization to improve therapy efficacy predictability.

We propose use of PDX therapy trials in preclinical testing of therapy strategies being considered for clinical translation as an effective mechanism to address the profound inter-tumoral heterogeneity that serves as a hurdle in therapy durability. In contrast to *in vitro* studies that observe significant benefit to targeting the PI3K/AKT/mTORC1 axis or the c-MET RTK to address BRAFi/MEKi resistance, our R-PDX *in vivo* therapy trials recapitulate clinical trials findings demonstrating a lack of efficacy from PI3K, mTOR, and cMET inhibitors in combination with BRAFi/MEKi in melanoma patients. We propose leveraging this melanoma R-PDX paradigm for preclinically screening of potential therapy regimens to improve likelihood of robust efficacy in phase 1 and phase 2 clinical trials. Our data demonstrate a relationship between pan-RTK protein expression and sensitivity to pan-RTK inhibitors. In addition, our R-PDX repurposing screen identifies the combination of dasatinib + BRAFi/MEKi has having the greatest potential as a salvage therapy strategy for melanoma patients that have already relapsed to standard of care therapy. Proteomic characterization of sensitive versus insensitive R-PDX models to the dasatinib combination identifies the expression of stem-like features in melanomas most likely to respond, which are historically difficult to therapeutically address.

## METHODS

Detailed standard operating procedures for all aspects of PDX generation and use are provided here (26, 27). Patient samples were collected under IRB approval.

### PDX Maintenance

All mouse therapy trials were performed in accordance with Wistar IACUC approved institutional guidelines. R-PDX were expanded in NSG mice until tumors reached 100-200 mm^3^. PDX bearing mice were than randomized into treatment groups, with 3 mice per group. Tumor size was measured three times a week using calipers, and tumor volume was estimated using the following formula; (length x width x width) / 2. Mice were sacrificed after 3-6 weeks of treatment.

### Massively Parallel Sequencing

DNA from PDX were characterized by massively parallel sequencing using a custom-designed 108-gene targeted seq panel (26, 28). Results were annotated for mutations and copy number changes. To address potential mouse DNA contamination, previously unreported variants with an allelic fraction of <0.15 were filtered out (26).

### scRNA-Seq methods

CellRanger suite (v3.1.0, https://support.10xgenomics.com) was used for pre-processing the scRNA-Seq data with Homo sapiens version GRCh38.93 as the reference transcriptome for aligning reads using STAR (29). Pre-filtering of low-quality cells was done by discarding cells with less than 200 genes with reads as well as those with over 10% mitochondrial content. This resulted in 8556 cells in DT and 5557 cells in DT+D. Sample integration was performed to correct for batch effect using the R package Seurat v4 (30). Seurat was also used to perform cell clustering, marker identification, and visualization. Differentially expressed genes between DT+D vs DT in each Seurat cluster were identified using Wilcoxon rank sum test and p values were adjusted with Bonferroni correction. Density plots of several known gene markers were used to identify cell subpopulations of interest.

### RPPA

Tumor lysates were prepared as previously described (31) and RPPA was performed by the MD Anderson Center RPPA core facility (Houston, TX, USA).

## Acknowledgments

The research was supported by NIH grants R01 CA 182890, U54 CA224070, P01 CA114046, P01 CA025874, P30 CA010815, R01 CA047159, K01 CA245124-01, the Dr. Miriam and Sheldon G. Adelson Medical Research Foundation, and the Melanoma Research Foundation. The support for Shared Resources utilized in this study was provided by Cancer Center Support Grant (CCSG) CA010815 and S10 OD023586 to the Wistar Institute.

## FIGURE LEGENDS

**Supplemental Figure 1:**
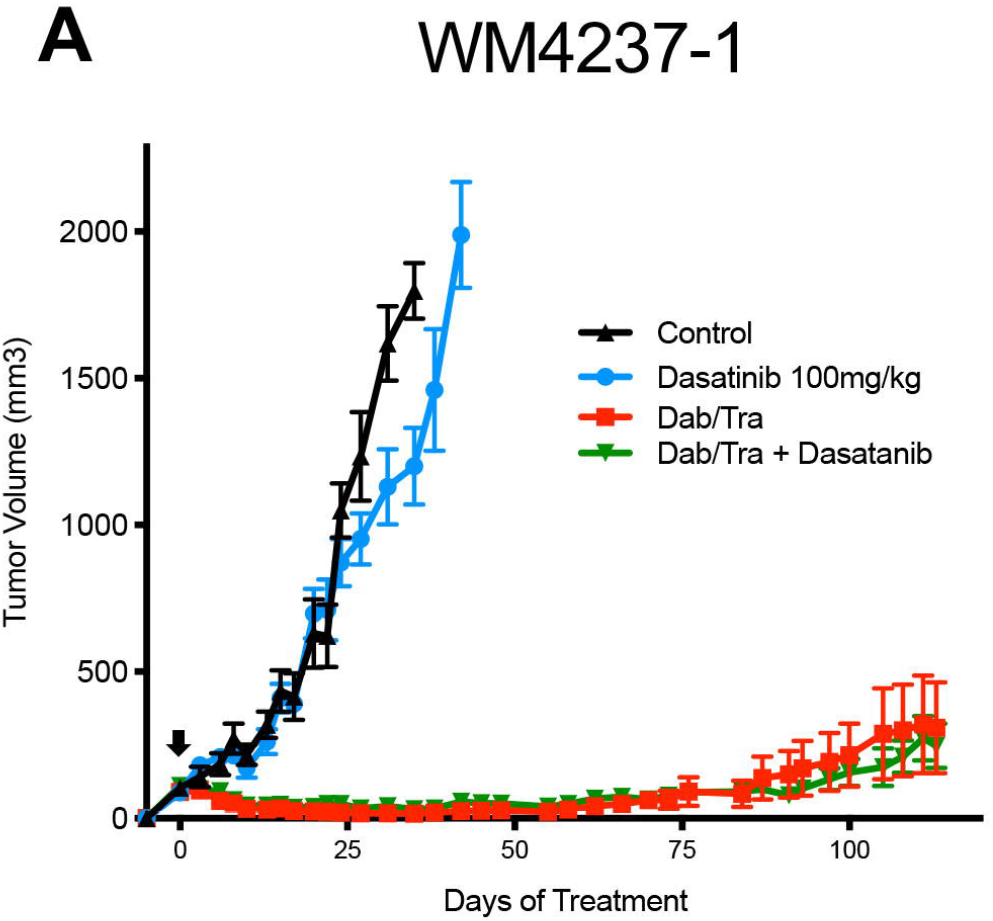
Dasatinib does not enhance BRAFi/MEKi efficacy in therapy naïve setting. (**A**) The therapy naïve PDX model WM4237-1 was implanted s.q. into an NSG mouse. Treatment with vehicle control, dasatinib, BRAFi//MEKi, or dasatinib + BRAFi/MEKi began when tumors were palpable (denoted with black arrow).

